# Hunters or gardeners? Linking community structure and function of trap-associated microbes to the nutrient acquisition strategy of a carnivorous plant

**DOI:** 10.1101/197020

**Authors:** Dagmara Sirová, Jiří Bárta, Karel Šimek, Thomas Posch, Jiří Pech, James Stone, Jakub Borovec, Lubomír Adamec, Jaroslav Vrba

**Affiliations:** Biology Centre CAS, Institute of Hydrobiology, Na Sádkách 7, CZ-37005, Èeské Budìjovice, Czech Republic; Faculty of Biological Sciences, University of South Bohemia, Branišovská 1760, CZ-37005 Èeské Budìjovice, Czech Republic; Limnological Station, Department of Plant and Microbial Biology, University of Zurich, Kilchberg CH-8802, Switzerland; University of Alaska Fairbanks, Department of Biology and Wildlife, Fairbanks, AK-99775, USA; Institute of Experimental Botany CAS, Rozvojová 263, CZ-16502, Praha 6-Lysolaje, Czech Republic; Institute of Botany CAS, Section of Plant Ecology, Dukelská 135, CZ-37982 Třeboň, Czech Republic

**Keywords:** algae, bacteria, ciliate bacterivory, digestive mutualism, fungi, herbivory, nutrient turnover, plant–microbe interactions, protists, *Utricularia* traps

## Abstract

All higher eukaryotes live in a relationship with diverse microorganisms which colonize their bodily surfaces; plants are no exception. However, we still lack a satisfactory understanding of how these loosely associated microbiomes with immense diversity and functional potential interact with their hosts or how these interactions shape processes within populations and ecosystems. There is considerable similarity between microbial communities colonizing plant surfaces such as roots, and those of the animal gut. This often overlooked parallel allows us to look at microbial as well as host ecophysiology from a fresh perspective. The traps of carnivorous plants are sophisticated digestive organs and interface environments between the supply and the demand for nutrients. We selected the miniature ecosystem in the traps of aquatic carnivorous *Utricularia* plants as our model system. By assessing the trap-associated microbial community structure, diversity, function, as well as the nutrient recycling potential of bacterivory, we gained insight into the nutrient acquisition strategies of the *Utricularia* hosts. We conclude that trap ecophysiological function is in many aspects highly analogous to that of the herbivore gut and centers around complex microbial consortia, which act synergistically to covert complex organic matter, often of algal origin, into a source of nutrients for the plants.

## Introduction

Recent progress in our understanding of the interdependencies between complex multicellular organisms and their associated microbes has led to an increasing awareness that these interactions are neither rare nor specialized. Instead, they represent fundamentally important aspects of biology and ecology (McFall-Ngai *et al.*, 2013).

Plant-associated microorganisms have long been recognized as key partners in enhancing plant nutrient acquisition, mitigating plant stress, promoting growth, or facilitating successful defense mechanisms against pathogens or grazers (for review, see Berg *et al*., 2014). Apart from the well-studied and relatively “simple” symbioses such as mycorrhizal and rhizobial interactions, there is a large pool of diverse microorganisms in varying degrees of association to different plant surfaces and tissues. These often highly complex microbial communities clearly play a significant role in plant ecophysiology, but the underlying mechanisms governing these associations are largely unexplored (Rinke *et al.*, 2013).The liquid-filled traps of carnivorous plants and phytotelmata of other species, such as tank-bromeliads, represent an environment with conditions highly suitable for microbial colonization and growth. They have often been used in the past as model systems for ecological research, especially focused on microbial food-web structure and related ecological questions (Cochran-Stafira and Von Ende, 1998; Koopman *et al.*, 2010; Gray *et al.*, 2012). However, the information on the relationship between microbial communities associated with these environments and their plant hosts has been scarce (Adlassnig *et al.*, 2011).

Recently, the hypothesis that microbial communities inhabiting the traps of rootless aquatic carnivorous *Utricularia* plants may be closely coupled to plant metabolism, nutrient acquisition, and growth was proposed (Sirová *et al.*, 2009). The exudation of large amounts of bioavailable photosynthates into *Utricularia* traps and their subsequent rapid utilization by the microorganisms present has been experimentally confirmed and represents a direct link between the plant host and associated microbiota (Sirová *et al.*, 2010, 2011). *Utricularia* are among the most numerous and cosmopolitan genera of carnivorous plants, attractive to researchers, due to their extremely small and unusual genomes (e.g., Carretero-Paulet *et al*., 2015; Silva *et al*., 2016). Depending on the species and growth conditions, a single *Utricularia* plant may bear hundreds to thousands of traps, usually on highly segmented leaves (Figure 1). These are tiny (1 – 5 mm long) liquid-filled bladders, whose lumen is isolated from the ambient environment by a two-cell thick trap wall and is generally anoxic or anaerobic (Adamec, 2007). The presence of various microbial taxa actively colonizing the lumen of *Utricularia* traps has previously been detected and roughly quantified (Sirová *et al.*, 2009, 2011; Caravieri *et al.*, 2014, Alcaraz *et al*., 2016). Likewise, it has been noted that the external surfaces of *Utricularia* plants tend to be covered in thick algal periphyton, exceeding in abundance that of other aquatic species (Díaz-Olarte *et al.*, 2007). Although *Utricularia -* associated microorganisms are thought to play an important role in the plant’s ecophysiology, a deeper insight into their community structure and ecology is lacking.

**Figure 1.**
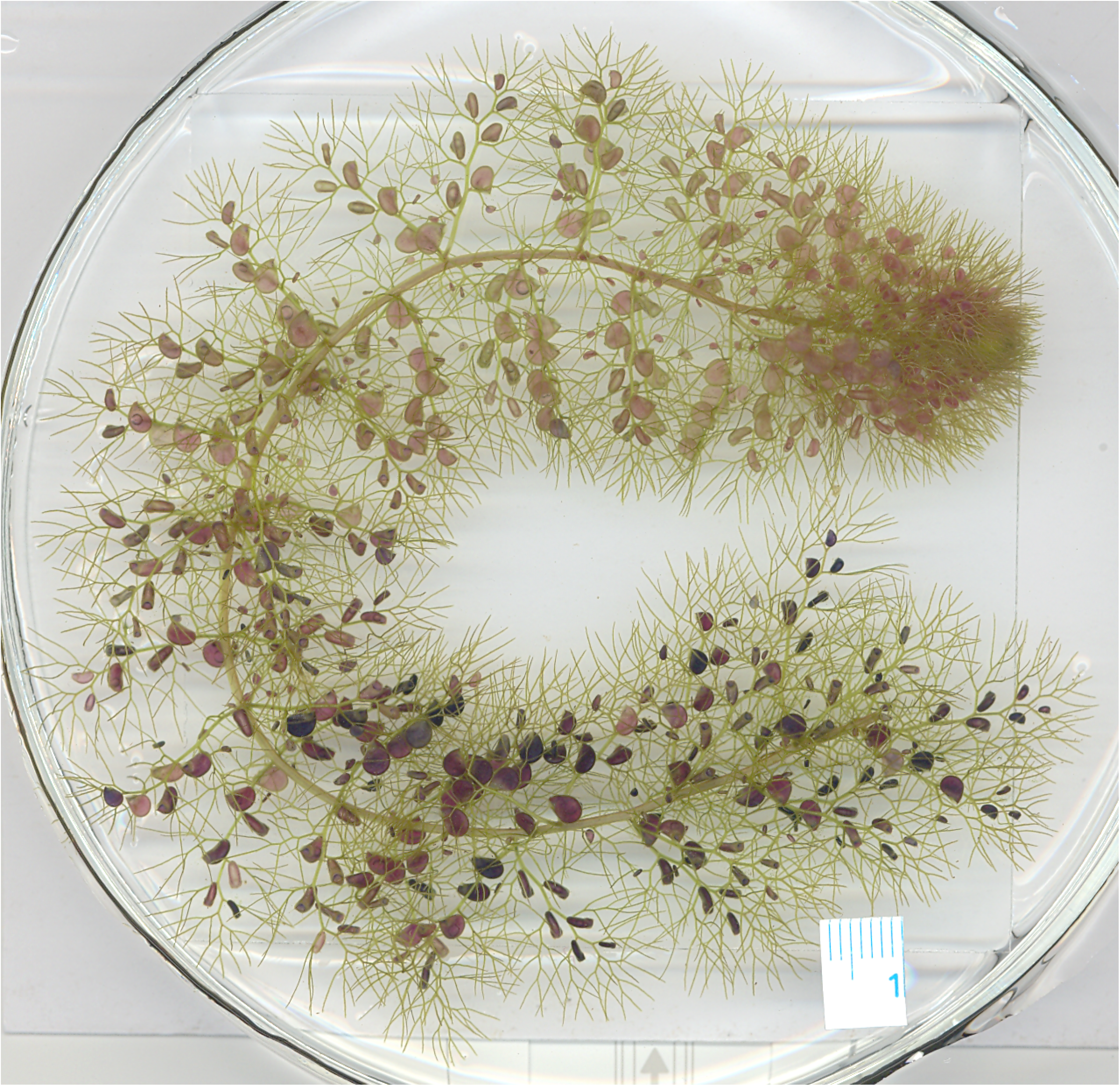
Experimental *Utricularia vulgaris* shoot on a Petri dish. Segmented leaves bearing traps and the growth tip are visible.

*Utricularia* were long thought to be classical examples of the carnivorous habit (Darwin, 1875). However, their nutrient acquisition strategy is the subject of debate and the importance of carnivory in their nutrition has been questioned (Richards, 2001; Sirová *et al.*, 2009). It has been proposed that algae, frequently observed and abundant in both the plant periphyton and traps (Sirová *et al.*, 2009; Plachno *et al.*, 2012), rather than metazoan plankton, are the main source of nutrients for the plants (Peroutka *et al.*, 2008; Alkhalaf *et al.*, 2009).

Here, we present the results of next generation sequencing of the microbiome, including the periphytic communities, associated with two *Utricularia* species – *U. australis* and *U. vulgaris*. Microbial community structure, diversity, the nutrient recycling potential by bacterivorous protists as well as their importance for plant growth are assessed in *U. reflexa*. It has been well established in both vertebrates and invertebrates that microbial diversity in the digestive tract reflects closely the composition of the prevailing food source (Nalepa *et al.*, 2001; Barboza *et al.*, 2010). The traps of *Utricularia* species also function as digestive systems and are interfaces between the “food” supply and the demand for nutrients of the rootless plants. The main nutrient source in *Utricularia*, however, has remained ambiguous and its direct verification at *in-situ* conditions technically challenging (Adamec, 2011). During trap lifespan, only a minority of traps capture a macroscopic prey, while all of them contain communities of microbial commensals (Friday, 1989; Richards, 2001).

Based on the observations of trap-associated microbial community structure and behavior and reports from literature, we hypothesize here that aquatic *Utricularia* traps are structures able to process and utilize organic material of algal (and/or plant) origin. To test this hypothesis, we assessed the functional capabilities of trap-associated microorganisms with respect to gene expression, and compared the microbial community structure with similar datasets from pitcher traps of other carnivorous plants, gut microflora of various vertebrates and invertebrates, including carnivores and detritivores, the rumen microbiota of various herbivores, and environmental samples from the soil, rhizosphere, and freshwater. By combining molecular methods, microscopy, and other approaches, we were able to shed light onto the ecology of this highly specific and unique microbial niche which, surprisingly, seems to bear closest analogy to mammalian rumen ecosystems.

## Material and Methods

### Plant material and sampling

To assess the presence and diversity of trap associated microbiota, the ecophysiologically well characterized aquatic species *U. vulgaris* and *U. australis* were selected. Adult *U. australis* plants (40 – 50 cm in length) were collected from a mesotrophic bog at Ruda fishpond (South Bohemia, see Sirová *et al.*, 2010 for details). The 0.8 m^2^ polypropylene experimental container, where *U. vulgaris* plants were cultivated, contained *Carex* sp. litter and approximately 250 liters of dystrophic water, closely simulating natural conditions (Adamec *et al.*, 2010a; Borovec *et al.*, 2012). For the assessment of microbial community structure, *U. vulgaris* and *U. australis* plants were divided into three sections of increasing age (young, mature, old). From each of these segments approximately 70 larger, excised, functional traps without visible metazoan prey and trapless leaves with periphyton were collected (approximately 200 mg FW).

To assess the actively transcribed microbial gene pool, excised *U. vulgaris* traps from the entire shoot without visible metazoan prey were collected randomly (approximately 250 mg FW), with pooled traps from a single plant considered a replicate; three biological replicates were collected in total. All collected plant material was immediately placed into liquid N_2_ and samples were stored at – 80°C until further processing.

For the estimation of the protozoan grazing rates, a different species with larger traps – the tropical *U. reflexa* – was selected, because relatively large volumes of trap fluid are needed for this analysis. The plants were cultivated indoors in 3-liter aquaria; in dystrophic cultivation water closely simulating natural conditions (see Adamec 2007).

### *SSU rRNA amplicon sequencing for the taxonomical evaluation of the* Utricularia*-associated microorganisms*

Nucleic acid extractions were conducted according to a modified bead-beating protocol (Urich *et al.*, 2008). Total DNA was quantified fluorometrically using SybrGreen (Leininger *et al.*, 2006). The PCR primers (515F/806R) targeted the V4 region of the SSU rRNA, previously shown to yield accurate phylogenetic information and to have only few biases against any bacterial taxa (Liu *et al.*, 2007; Bates *et al.*, 2011; Bergmann *et al.*, 2011). Each sample was amplified in triplicate, combined, quantified using Invitrogen PicoGreen and a plate reader, and equal amounts of DNA from each amplicon were pooled into a single 1.5-ml microcentrifuge tube. Once pooled, amplicons were cleaned up using the Ultra Clean PCR clean up kit (MO BIO Laboratories). Amplicons were sequenced on the Illumina MiSeq platform at the Institute of Genomics and Systems Biology, Argonne National Laboratory, Argonne (Chicago, USA). Paired-end reads were joined using Perl scripts yielding approximately 253bp amplicons. Approximately 1.8 million paired-end reads were obtained with average 66.000 reads per sample. Quality filtering of reads was applied as described previously (Caporaso *et al.*, 2010). Reads were assigned to operational taxonomic units (OTUs, cut-off 97% sequence identity) using an open-reference OTU picking protocol with QIIME implemented scripts (Caporaso *et al.*, 2010). Reads were taxonomically assigned using Green Genes database, release 13.08 as reference. Those reads which were assigned as “chloroplast” and “mitochondrion” were excluded from further analyses.

### Meta-analyses of prokaryotic communities from different habitats

To compare the composition of *Utricularia* trap-associated microbial communities, data (OTU tables in biom format) from 9 relevant studies represent different habitats were analyzed. Seven of the studies were downloaded from Qiita (https://qiita.ucsd.edu/) database, 16S rDNA sequences of the *Nepenthes* pitcher microbiome from the SRA archive (project ID PRJNA225539) and *Sarracenia* pitcher sequences from the Genebank database (accession numbers JF745346 – JF745532 and JN368236 – JN368422) (Supplementary Table 1). Altogether, 4221 samples were included in the meta-analysis. The Qiita database works with the closed reference OTU picking algorithm, we have therefore processed our sequences and also the sequences from *Nepenthes* and *Sarracenia* pither microbiomes in the same way as the Qiita pipeline, to ensure comparability with Qiita OTU tables. All OTU tables were then merged together using Qiime scripts and analyzed as one dataset. Non-metric multidimensional scaling (NMDS) using Bray-Curtis dissimilatory metrics was used for computing distances between samples. Because the large amount of samples reduced clarity of the graphical representation of results, for illustrative purposes only, we randomly chose 30 samples from each study to construct a dendrogram (Figure 2a) using the UPGMA algorithm.

**Figure 2.**
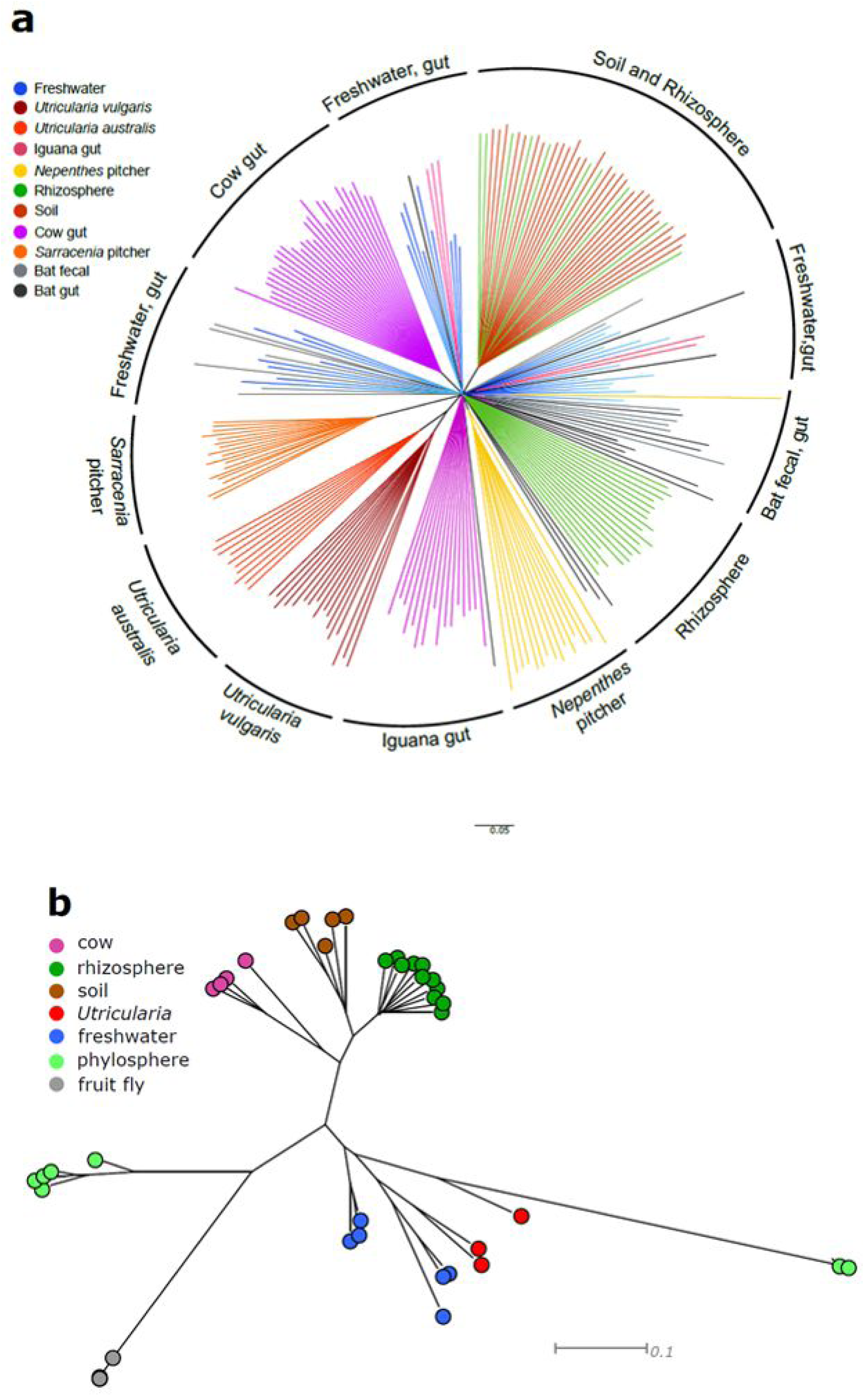
Meta-analyses of prokaryotic communities from different environments (a) based on 16S rDNA gene sequencing (4421 samples, UPGMA tree, data from 9 relevant studies were downloaded from the Qiita database (https://quiita.ucsd.edu/) and metagenomes/metatranscriptomes (b) based on shotgun sequencing and KEGG annotations (43 samples obtained from MG-RAST, Neighbor joining tree). For better visualization of the UPGMA tree (b), only up to 30 samples per study were selected.

To obtain a more function-related point of view, three *Utricularia vulgaris* metatranscriptomes were compared to 43 metagenomes/metatranscriptomes from 6 different habitats. The sequences from these studies were obtained from the MG-RAST server (Table S1)

### *RNA-seq analysis for functional profiling of the* U. vulgaris *trap-associated microbiome*

To assess the actively transcribed microbial gene pool, total RNA was extracted from the *U. vulgaris* trap samples (n = 3), using the protocol identical to that described in detail previously (Sirová *et al.*, 2014). Briefly, DNA was removed from the extracts and two transcriptomic libraries, eukaryotic and prokaryotic, were created at the Institute of Genomics and Systems Biology, Argonne National Laboratory, Argonne (Chicago, USA) using standard Illumina TruSeq RNA library preparation kits. The ribosomal RNA as well as eukaryotic (plant) RNA fraction was removed in order to enrich prokaryotic transcripts, and, *vice versa*, eukaryotic transcript enrichment was performed in parallel, to capture transcripts from fungi, protists, and other eukaryotic microorganisms. Enriched mRNA from both libraries was reverse transcribed to create metatranscriptomic libraries and sequenced using Illumina HiSeq platform (100×100 cycle paired-end run). We obtained approximately 40 million paired-end reads per sample. Reads were quality checked, low quality reads and reads with ambiguous bases were filtered out. Reads from all three replicates (approx. 120 million sequences) were then assembled with Velvet Optimizer (Zerbino and Birney, 2008) which resulted in approximately 500,000 contigs. In this step, we filtered out the potential *Utricularia* transcripts by blasting contigs against our *Utricularia* reference transcriptome (Bárta et al. 2015). All contigs which gave significant hit (e-value <0.0001, min score 80) were excluded from further analyses. Contigs were also blasted against the SILVA database (release 111) to identify ribosomal RNAs (rRNAs). Those sequences that gave BLAST bit score greater than 80 were marked as potential rRNAs and extracted from the dataset. Reads were than mapped back onto the remaining contigs using bowtie algorithm with default parameters (Haas *et al*., 2013). To identify potential prokaryotic functional gene transcripts, the remaining contigs without rRNAs were blasted against the nr database with e-value of 0.001.

### Microbial network analyses

The relative abundances of OTUs were square-root transformed and the 300 most abundant OTUs in *U. vulgaris* samples were selected, to avoid the spurious correlations of rare OTUs. The resulting OTU tables, separately for the trap (15 samples) and the periphyton (16 samples) were used for microbial network analyses. The analysis was done in Cytoscape 3.0.2 with the CoNET application (Shannon *et al.*, 2003; Faust *et al.*, 2012). The parameters and settings for network analyses in CoNET application were:-parent_child_exclusion,-row_minocc 8,-correlations (Spearman, Pearson, mutual information, Bray – Curtis and Kullback – Leibler dissimilatory). The threshold for edge selection was set to 1,000 top and bottom. During randomization, 100 iterations were calculated for edge scores. In the following bootstrap step, 100 iterations were calculated, unstable edges were filtered out. The Brown method was chosen as the P-value merging method and the Benjamini – Hochberg procedure was selected for multiple test correction. The resulting network was visualized and analyzed (i.e. degree of nodes, betweenness centrality, closeness centrality) in Cytoscape 3.0.2. Potential keystone OTUs were identified (Berry and Widder, 2014).

For the estimation of unique microbiomes of *U. australis* and *U. vulgaris* traps and periphyton, the OTUs were grouped at the genus level. Genera presented only in *U. australis* or *U. vulgaris* and respective trap or periphyton samples were classified as unique microbiomes.

### Quantification of trap-associated eubacterial and fungal communities

Quantification of bacterial and fungal SSU rRNA genes was performed using the FastStart SybrGREEN Roche^®^ Supermix and Step One system (Life Technologies, USA). Each reaction mixture (20 μl) contained 2 μl DNA template (∽1 – 2 ng DNA), 1 μl each primer (0.5 pmol μl^−1^ each, final concentration), 6 μl dH_2_O, 10 μl FastStart SybrGREEN Roche^®^ Supermix (Roche, France) and 1 μl BSA (Fermentas, 20 μg μl^-^1). The qPCR conditions for bacterial quantification were as follows: initial denaturation (3 min, 95°C) was followed by 30 cycles of 30 s at 95°C, 30 s at 62°C, 15 s at 72°C, and completed by fluorescence data acquisition at 80°C used for target quantification. Product specificity was confirmed by melting point analysis (52°C to 95°C with a plate read every 0.5°C) and amplicon size was verified with agarose gel electrophoresis.

Bacterial DNA standards consisted of a dilution series (ranging from 10^1^ to 10^9^ gene copies μl^−1^) of a known amount of purified PCR product obtained from genomic *Escherichia coli ATCC 9637* DNA by using the SSU gene-specific primers 341F/534R (Muyzer *et al.*, 1993). The R^2^ values for the standard curves were >0.99. Slope values were >3.37 giving an estimated amplification efficiency of >93%. The qPCR conditions for fungal quantification were as follows: initial denaturation (10 min, 95°C) followed by 40 cycles of 1 min at 95°C, 1 min at 56°C, 1 min at 72°C, and completed by fluorescence data acquisition at 72°C used for target quantification. Fungal DNA standards consisted of a dilution series (ranging from 10^1^ to 10^7^ gene copies μl^-^1) of a known amount of purified PCR product obtained from genomic *Aspergillus niger* DNA by using the SSU gene-specific primers nu-SSU-0817-5’ and nu-SSU1196-3’ (Borneman and Hartin, 2000). R^2^ values for the fungal standard curves were > 0.99. The slope was between −3.34 to −3.53 giving estimated amplification efficiency between 95 and 93%, respectively.

Detection limits for the various assays (i.e. lowest standard concentration that is significantly different from the non-template controls) were less than 100 gene copies for each of the genes per assay. Samples, standards, and non-template controls were run in duplicates. To deal with potential inhibition during PCR, the enhancers (BSA, DMSO) were added to the PCR mixture. Also several dilutions (10×, 20×, 50×, 100×, and 1000×) for each sample were tested to see the dilution effect on Ct values.

### Quantification of trap-associated methanogenic and methanotrophic communities

Quantification of the *mcrA* gene (methanogens) was performed using the FastStart SybrGREEN Roche^®^ Supermix and Step One system (Life Technologies, USA). Each reaction mixture (20 μl) contained 2 μl DNA template (∽1 – 2 ng DNA), 0.1 μl each primer (0.3 pmol μl^−1^ each, final concentration), 6 μl dH_2_O, 10 μl FastStart SybrGREEN Roche^®^ Supermix (Roche, France) and 0.4 μl BSA (Fermentas, 20 μg μl^−1^). Primers ME1 (5’-GCM ATG CAR ATH GGW ATG TC-3’) a MCR1R (5’-ARC CAD ATY TGR TCR TA-3’) producing amplicon length of 280bp (Hales *et al*., 1996). The qPCR conditions for *mcrA* gene were as follows: initial denaturation (10 min, 95°C) followed by 35 cycles of 30s at 95°C, 1 min at 50°C, 1 min at 72°C, and completed by fluorescence data acquisition at 72°C used for target quantification. Standards consisted of a dilution series (ranging from 10^1^ to 10^7^ gene copies μl^−1^) of a known amount of purified PCR product obtained from genomic DNA of *Methanosarcina barkeri* DSM-800.

Quantification of *pmoA* gene (methanotrophs) was performed using the FastStart SybrGREEN Roche® Supermix and Step One system (Life Technologies, USA). Each reaction mixture (20 μl) contained 2 μl DNA template (∽1 – 2 ng DNA), 0.24 μl each primer (0.5 pmol μl^−1^ each, final concentration), 5.6 μl dH_2_O, 10 μl FastStart SybrGREEN Roche® Supermix (Roche, France), 0.5 μl DMSO and 0.4 μl BSA (Fermentas, 20 μg μl^−1^). Primers A189-F (5’-GGNGACTGGGACTTCTGG-3’) a M661-R (5’-GGTAARGACGTTGCNCCGG-3’) producing amplicon length of 491bp (Kolb *et al*., 2003). The qPCR conditions for *pmoA* gene were as follows: initial denaturation (10 min, 95°C) followed by 35 cycles of 30 s at 95°C, 20 s at 57°C, 45 s at 72°C, and completed by fluorescence data acquisition at 72°C used for target quantification. Standards consisted of a dilution series (ranging from 10^1^ to 10^7^ gene copies μl^-^1) of a known amount of purified PCR product obtained from genomic DNA of *Methylobacter luteus*.

### Sequence deposition

Raw sequences of 16S rDNA, ITS1 amplicons, and raw sequences of all three metatranscriptomes were deposited in European Nucleotide Archive (ENA) under study ID PRJEB19666. Annotated metatranscriptomic sequences can be found at the following website: <u>http://utricularia.prf.jcu.cz/</u>

### Bacterial and protozoan quantification and the estimation of protozoan grazing rates

Ten *U. reflexa* plants were divided into three segments of increasing age (young, mature, old). Each segment contained 6 leaf whorls. Trap fluid was collected from the traps in each segment (see Sirová *et al.*, 2009) and a pooled sample (∽750 μl for each age category) from all ten plants was made. Bacterial and protozoan counts in the trap fluid samples were estimated using epifluorescence microscopy, according to methods described previously (Šimek *et al.*, 2000).

The protist grazing rates were estimated using fluorescently labelled bacteria (FLB) as a tracer. The FLB were prepared from the strain *Limnohabitans planktonicus*, as detailed in Šimek *et al*. (2017). Cell sizes of the strain are comparable to that of bacterial cells commonly occurring within the *U. reflexa* traps. The FLB uptake rates were determined in short-term triplicate experiments, where tracer FLB were added to the trap-fluid samples to constitute 6-8% of the total bacterial concentration. For further details on sample fixation, protist staining and enumeration, and tracer ingestion determinations, see Šimek e*t al.* (2017). At least 45 ciliates were inspected for FLB ingestion in each replicate sample. To estimate total protist grazing, we multiplied average uptake rates of protozoa by their *in-situ* abundances as previously described (Šimek *et al.*, 2000).

## Results and Discussion

### Microbial diversity

Our results show that *Utricularia* plants harbor microbial communities rich in both taxonomic composition and metabolic capabilities. We have identified over 4,500 distinct OTUs in the traps and periphyton (see the complete OTU list in Supplementary Table 2a, 2b). Microbial (prokaryotic) alpha diversity in two studied *Utricularia* species was significantly higher than that found in the traps of other carnivorous plants and was comparable to that observed in other high-diversity systems: the rhizosphere and the cow gut (Table 1). A comparison with datasets from other environments with regards to composition (Figure 2a) revealed that the *Utricularia* trap microbiomes are species-specific, but similar between the two studied species. Out of the different environments selected, they most closely resembled the communities inhabiting the pitchers of a terrestrial carnivorous plant, *Sarracenia*, and those found inside the guts of herbivorous iguanas. When comparing the metatranscriptomic data from the traps of *U. vulgaris* with other systems, the highest similarity (more function-related) was with freshwater microbial communities (Figure 2b).

**Table 1.** Comparison of selected alpha diversity indexes for various 16S datasets from different habitats (*N =* number of samples in study). All datasets were subsampled to 2000 sequences prior to analyses, more information on the data used can be found in Supplementary Table 1.

| **Habitat** | **N** | **PD whole tree** | **ANOVA** | **Chao1 index** | **ANOVA** | **Obs. sp.** | **ANOVA** | **Shannon index** | **ANOVA** |
| --- | --- | --- | --- | --- | --- | --- | --- | --- | --- |
| Soil | 309 | 74.2 | a | 2238.5 | a | 1043.2 | a | 9.6 | a |
| Rhizosphere | 751 | 49.6 | b | 1505.3 | b | 675.5 | b | 7.3 | b |
| Cow gut | 188 | 35.0 | c | 778.5 | c | 480.1 | c | 7.6 | b |
| *Utricularia* | 22 | 34.2 | c | 816.4 | c | 422.5 | c | 7.0 | b |
| Trap | 16 | 35.0 | c | 859.2 | c | 439.3 | c | 7.1 | b |
| Periphyton | 6 | 32.2 | c | 702.2 | c | 377.7 | c | 6.9 | b |
| Iguana gut | 91 | 22.6 | b | 359.3 | d | 218.3 | d | 5.4 | c |
| Freshwater | 2635 | 21.1 | b | 262.9 | d | 206.9 | d | 5.4 | c |
| *Sarracenia*<br>pitchers | 17 | 15.4 | bd | 270.1 | d | 169.4 | de | 5.3 | c |
| *Nepenthes* pitchers | 16 | 13.1 | bd | 211.2 | d | 132.8 | de | 4.5 | c |
| Bat gut | 51 | 11.9 | d | 185.4 | d | 108.4 | e | 3.0 | de |
| Bat fecal | 13 | 10.3 | bd | 163.4 | d | 87.8 | de | 2.6 | e |

When comparing the trap and periphytic communities associated with one *Utricularia* species, the bacterial communities in both environments were dominated by Proteobacteria (58% and 54% of assigned sequences in periphyton and trap, respectively; p = 0.013) and both had very similar bacterial composition at the family level. Over half of the taxa were shared even at the genus level (Figure 3). The Comamonadaceae were the most abundant group, constituting similar proportion (>10%) of both the trap and the periphytic communities. The close similarity of periphytic and trap communities suggests strong links between *Utricularia* external surfaces and the internal trap environment, with periphyton being the most likely source of inoculum for microbial colonization of traps (Sirová *et al*., 2009; 2011).

**Figure 3.**
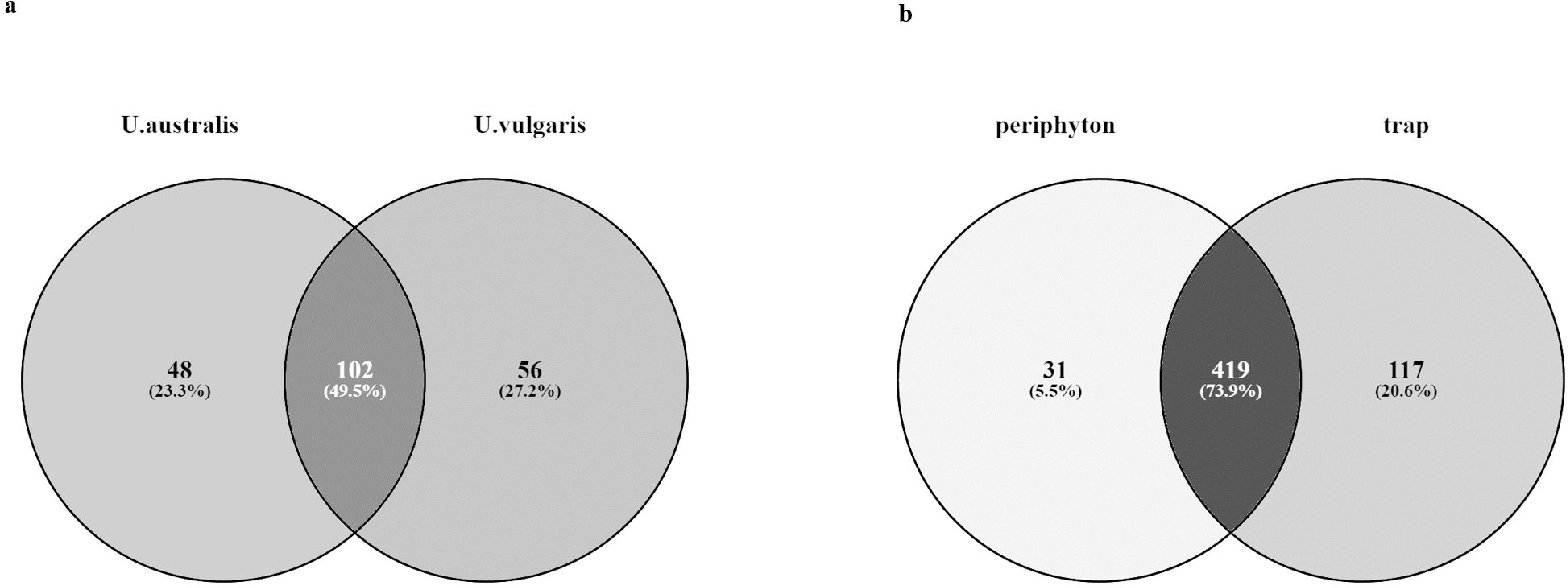
Compositional overlap in *Utricularia*-associated microbiomes at the genus level, computed for prokaryotic genera only. (a) Comparison between *U. australis* and *U.vulgaris* microbiomes and (b) between *U. australis* and *U.vulgaris* periphyton and trap.

Despite of the high similarity between the trap lumen and the periphyton in terms of prokaryotic community composition, when we compare the microbial co-occurrence analyses, we see two strikingly different systems with distinct “key-stone” taxa (Supplementary Table 3) and a distinct degree of interconnectedness (Supplementary Figures 1, 2) in each of the two environments. While the periphytic communities show a co-occurrence pattern (Supplementary Figure 1) typically observed in highly spatially and functionally interconnected microbial biofilms, the co-occurrence pattern of the trap community consists of several smaller, mutually disconnected microbial networks and implies a much more heterogeneous and fragmented environment inside the trap lumen (Supplementary Figure 2). This result is consistent with previously published observations showing progressing degree of microbial aggregation into flocks and multispecies biofilms with progressing *Utricularia* trap development (Sirová *et al*., 2009). In analogy, results coming from direct microscopic examinations of the rumen ecosystem and its contents (McAllister *et al*., 1994) show that microbial populations are largely attached to feed particles (>70% of total numbers, Russell *et al.*, 2009) and that bacteria from the ruminal fluid associate into micro-colonies forming multispecies biofilms. These metabolically related organisms then associate with their preferred substrates and produce the myriad of enzymes necessary for the digestion of chemically and structurally complex plant tissues, hence, creating a system of mutually disconnected micro-environments.

In terms of core microbiota, one of the most distinct groups, which were specific to the *Utricularia* trap environment only, were members of the Peptostreptococcaceae family (Clostridiales), which successfully colonized the lumen, but were rarely found in the periphyton (Supplementary Table 2a). The group represents anaerobic fermenters often associated with habitats such as animal guts and oral cavities, manure, soil, and sediments (Slobodkin, 2014). In the rumen ecosystem, *Peptostreptococcus* spp. produce high amounts of ammonia, but are not able to hydrolyze intact proteins and do not use carbohydrates as a carbon source. Thus, they occupy a niche of peptide-and amino-acid-degrading microorganisms and depend on proteolytic bacteria (Attwood *et al.*, 1998). The genes expressed in the traps that were assigned to this family, such as pyrroline-5-carboxylate reductase [EC:1.5.1.2] or threonine aldolase [EC:4.1.2.5] (Supplementary Table 4), are all involved in the metabolism of amino acids. In ruminants, as much as 90% of the amino acids reaching the small intestine are derived from ruminal microorganisms. The trap lumen contains very high concentrations of ammonium ions (Sirová *et al.*, 2014) and many bacteria able to hydrolyze proteins are present; it is therefore likely that a process highly similar to that in the rumen involving Peptostreptococcaceae, is taking place therein.

**Table 2.** Comparison of the abundance of total bacteria, methanothrophs, methanogens, fungi, and fungal to bacterial ratio (F/B) in the trap and periphyton of *Utricularia* species. Quatification of bacteria, fungi, methanotrophs and methanogens was done using the 16SrDNA, 18SrDNA, pmoA, and mcrA gene copy numbers, respectively. Quantity was normalized to total amount of DNA.

|  | **Bacteria**<br>(10<sup>6</sup>) | **Fungi**<br>(10<sup>3</sup>) | **Methanotrophs**<br>(10<sup>6</sup>) | **Methanogens** | **F/B** |
| --- | --- | --- | --- | --- | --- |
| **Trap** (n=16) | 1.6 | 7.7 | 0.34 | B.D. | 0.0048 |
| *St.dev* | *1.9* | *5.3* | *0.4* |  |  |
| **Periphyton** (n = 6) | 2.9 | 41 | 0.41 | B.D. | 0.0143 |
| *St.dev* | *4.3* | *79* | *0.21* |  |  |

B.D. – below detection

### Methane metabolism

Herbivore gut ecosystems generally tend to produce copious amounts of methane as a result of the fermentation activity by the strictly anaerobic methanogenic Archaea (Hackstein and Van Alen, 1996). Using gas chromatography, we have not detected the release of methane gas from the *Utricularia* traps (data not shown), and methanogens were not found in our trap fluid samples using the qPCR assay (Table 2). However, significant amounts of methanotrophs were found, constituting up to 40% of total prokaryotic community (Supplementary Table 2c). These included active obligate methanotrophs, for example from the genus *Methylococcus* (Gammaproteobacteria, Supplementary Table 2c). Moreover, methane metabolism was also found to be one of the most expressed prokaryotic modules (KEGG) in the trap fluid metatranscriptome from *U. vulgaris* (Table 2a). These somewhat paradoxical results may be explained by a recent discovery (Repeta *et al*., 2016) that the degradation of dissolved organic matter in the aquatic environment by commonly occurring bacteria, e.g. *Pseudomonas* spp., can, even in the presence of oxygen, result in the release of methane, ethylene and propylene.

We speculate that this process can explain the presence and activity of methanotrophs in the *Utricularia* traps, which may metabolize all of the produced methane, thus preventing its detection. This hypothesis, however, needs to be experimentally verified.

### Degradation of complex algal/plant polymers

We have detected the presence of bacterial genera in the traps, which are considered to include cellulolytic species capable of degrading complex organic material of plant (algal) origin (Supplementary Table 2a), for example *Clostridium*, *Ruminococcus*, *Caldicellulosiruptor*, *Butyrivibrio*, *Acetivibrio*, *Cellulomonas*, *Erwinia*, *Thermobifida*, *Fibrobacter*, *Cytophaga*, and *Sporocytophaga*. Also notable is the significant presence of active myxobacteria (*Cystobacter* spp.), which are known to include cellulolytic species and are frequently isolated from systems with high decomposing plant material content (Shimkets *et al.*, 2006).

Bacterial degradation of cellulose-rich material is more restricted to biomass containing low amounts of lignin, as bacteria are generally poor producers of lignanases. Plant and algal biomass produced in aquatic systems contains little amounts of lignin and is typically degraded by bacteria which are better adapted for the aquatic environment than fungi. Overall numbers of bacterial cells found in the *U. reflexa* trap fluid were similar to those reported previously (Sirová *et al*., 2009; 2011), being an order of magnitude higher in the old traps compared to the young and mature (Table 3). Bacterial degradation of cellulose-rich material is more restricted to biomass containing low amounts of lignin, as bacteria are generally poor producers of lignanases. Plant and algal biomass produced in aquatic systems contains little amounts of lignin and is typically thought to be degraded mostly by bacteria, which are better adapted for the aquatic environment than fungi.

**Table 3.** The estimates of bacterial and ciliate numbers, the individual grazing rate (IGR) and total grazing rate (TGR) of ciliates and the turnover rate of bacterial standing stock (Turnover) in *U.reflexa* traps of different age is presented. Means of 3 technical replicates are shown.

| **Trap age** | **Bacteria**<br>[10<sup>6</sup> ml<sup>-1</sup>] | **Protozoa**<br>[10<sup>3</sup> ml<sup>-1</sup>] | **IGR**<br>[bact. prot.<sup>-1</sup> h<sup>-1</sup>] | **TGR**<br>[10<sup>6</sup> bact. ml<sup>-1</sup> d<sup>-1</sup>] | **Turnover**<br>[day<sup>-1</sup>] |
| --- | --- | --- | --- | --- | --- |
| Young | 44.5 | 35.4 | 273.4 | 223.1 | 5.0 |
| Mature | 65.3 | 46.5 | 263.1 | 290.0 | 4.4 |
| Old | 266.9 | 50.8 | 342.1 | 411.8 | 1.5 |

**Table 4.**
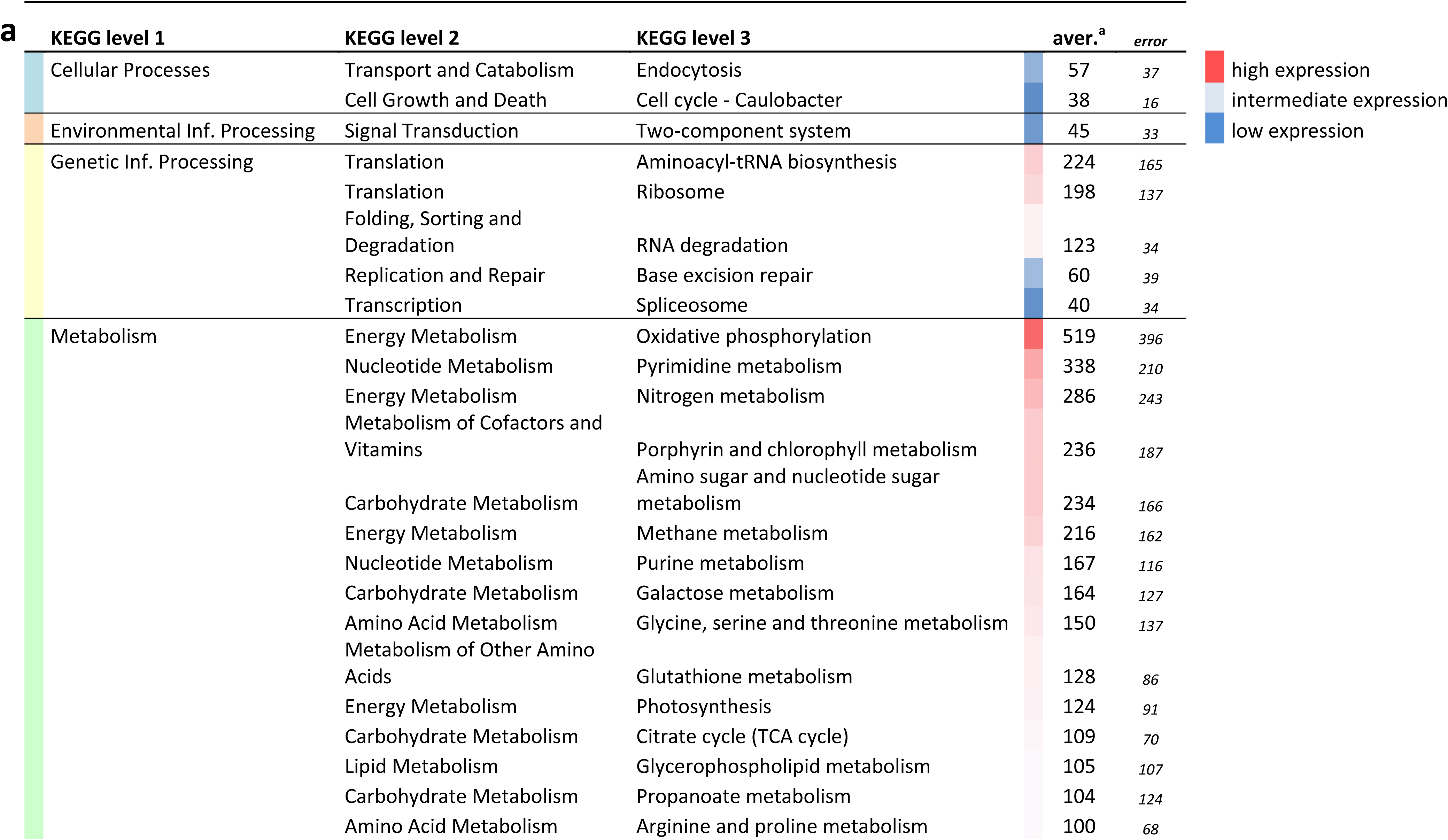
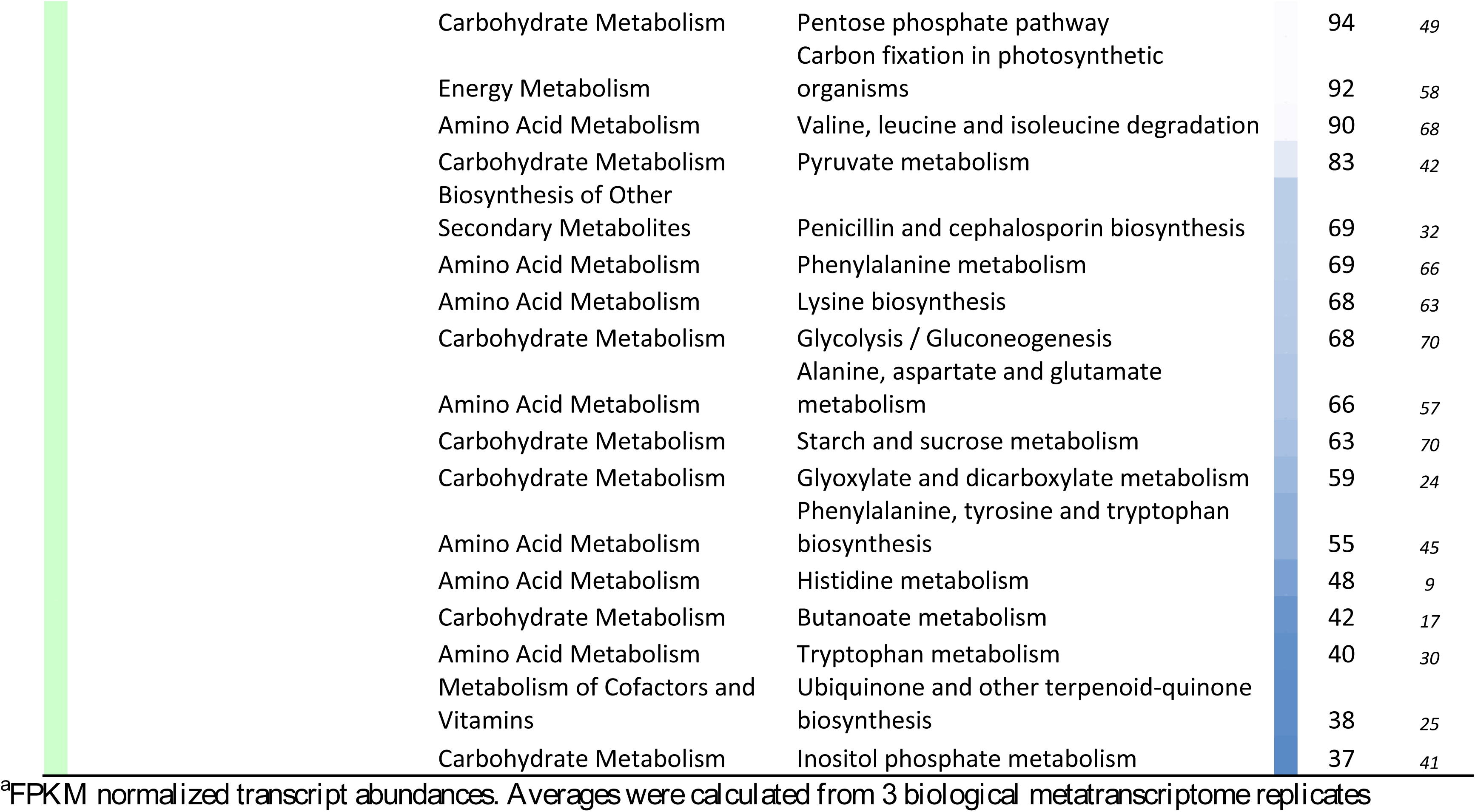
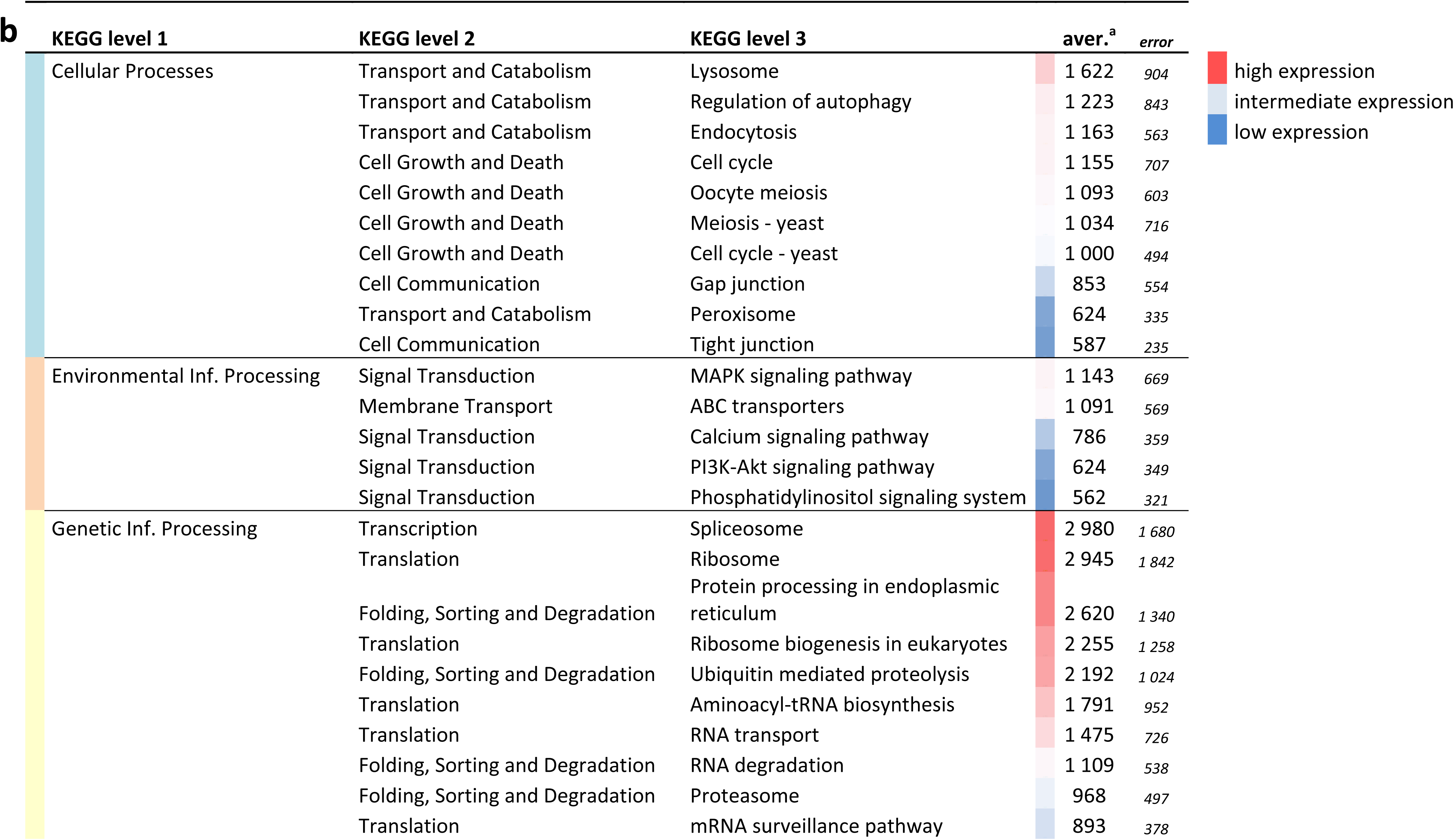
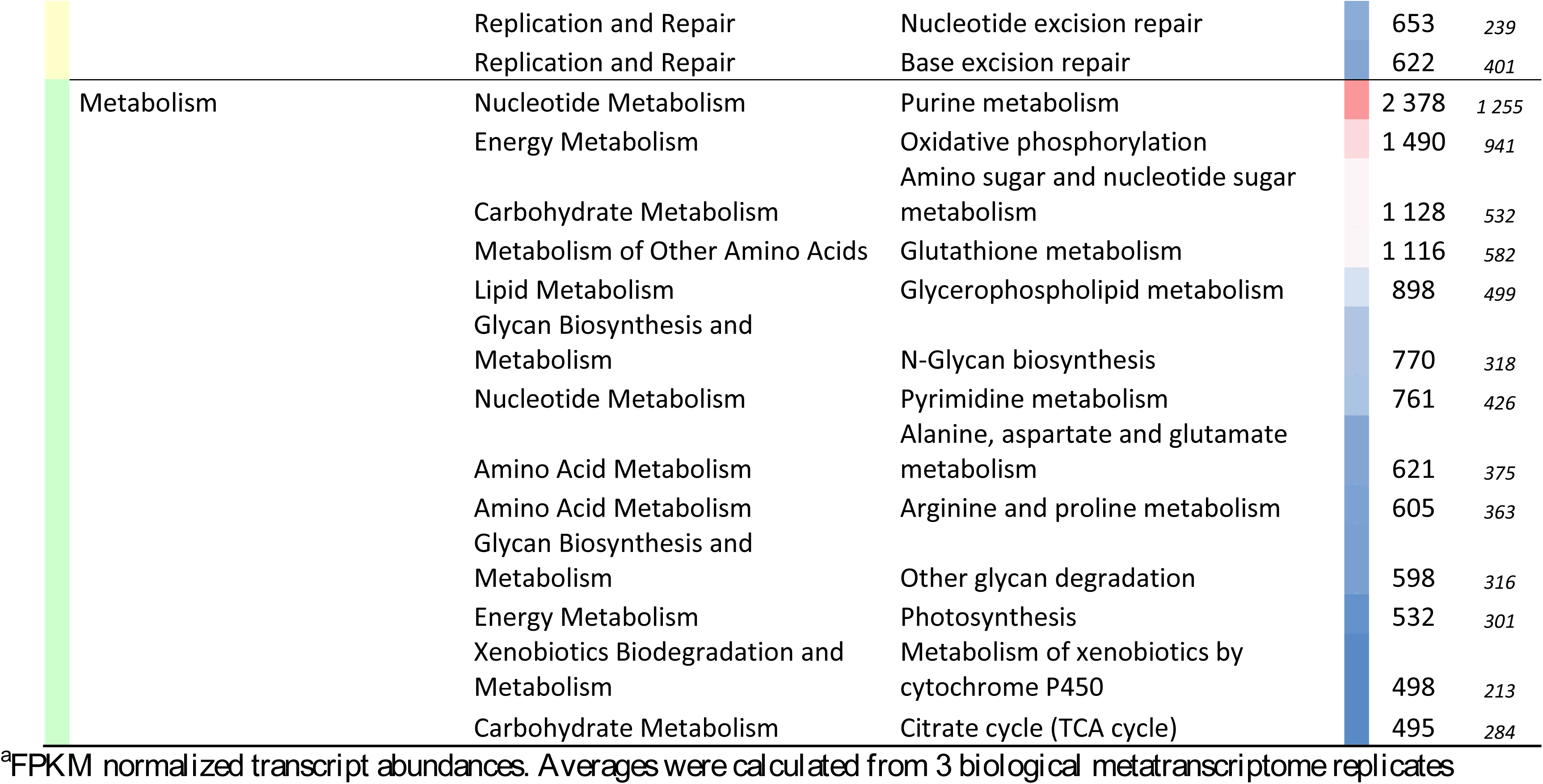
The 40 most expressed prokaryotic (a) and eukaryotic (b) functional KEGG modules. Functional assignment was done in MEGAN, using the KEGG database. Abundances were estimated in Trinity by mapping raw reads to assembled contigs, using the *bowtie2* algorithm

Fungi, however, were also found to be present in the traps and represented a significant proportion of the total microbial community, as quantified by qPCR (Table 2). Fungi, especially the anaerobic chytrids (Chytridiomycota), are frequently found in the gastrointestinal tracts of ruminants and non-ruminant herbivores, either associated with digested organic material, or as free-swimming zoospores in the fluid phase, playing significant role in the digestive process of complex plant polymers. Many fungal taxa whose presence inside of the *Utricularia* traps was determined by SSU rRNA sequencing, (e.g. *Chrysomphalina* sp., Agaricales, Basidiomycota, Supplementary Table 2b) were most probably entrapped as spores from the ambient environment and do not represent trap-associated microbes as such, but rather a potential source of nutrients. Others, most notably the saprotrophic *Basidiobolus* sp. (Basidiobolales, Zygomycota) abundant in the traps of all ages (up to 45% of total OTUs), or species belonging to the above mentioned Chytridiomycota (Supplementary Table 2b) found to be actively growing in the traps (Supplementary Table 4), are likely a component of the trap microbial network and contribute to the nutrient release and assimilation by the plant host.

pH is considered a key factor affecting the efficiency of organic matter degradation and nutrient release in the rumen and herbivore digestive tracts as both fungal and bacterial enzymes are sensitive to pH changes (Russel and Wilson, 1996). It has been shown that the trap pH in aquatic *Utricularia* is highly stable, usually acidic, species specific (Sirová *et al*., 2003, 2009) and its stability likely contributes to the optimal development of the trap-associated microbial communities and their activities.

### Protistan grazing and nutrient turnover

A third important group of microbes in the traps are the protists. Their numbers in the *U. reflexa* traps were extremely high, rising steadily with increasing trap age, up to approximately 50,000 cells per milliliter of trap fluid in the oldest traps. This means that a single trap, depending on age, harbored tens to hundreds of individuals (Table 3). Such high population densities are rarely found in natural environments and are comparable only to those found in the mammalian rumen or in the activated sludge systems (e.g. Towne *et al*., 1990; Abraham *et al*., 1997). The protist community in *U. reflexa* traps, however, had very low diversity and essentially consisted of a monoculture composed of various euglenid species and a single conspicuous zoochlorellae-bearing ciliate (Figure 4). This organism has recently been described as a new species – *Tetrahymena utriculariae* – and has not yet been found in any other environment (Pitsch *et al.*, 2017; Šimek e*t al.,* 2017).

**Figure 4.**
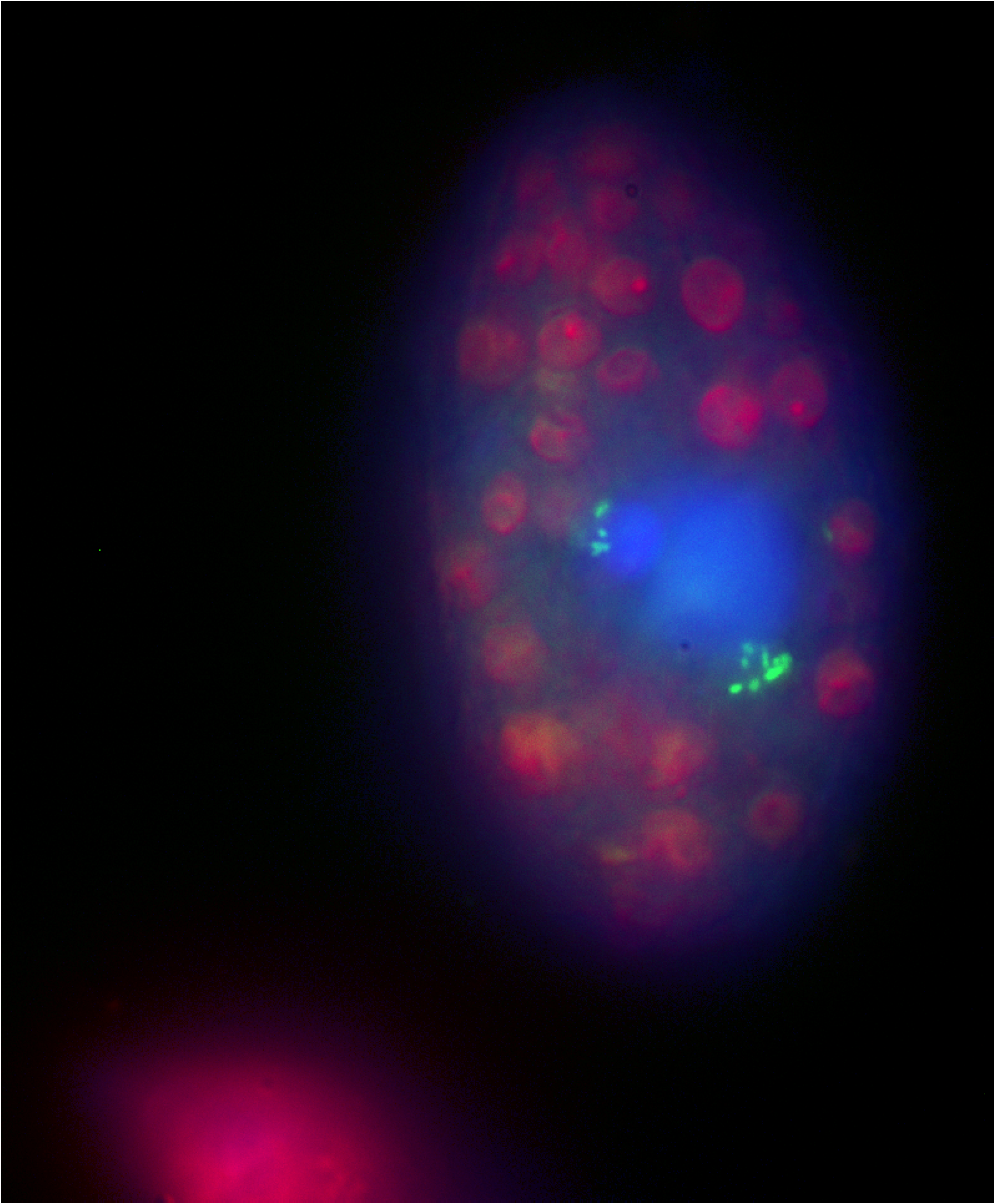
*Tetrahymena utriculariae* under the epifluorescence microscope. Zoochlorellae are visible in purple, the nucleus is stained blue, and fluorescently labeled bacteria in food vacuoles show green fluorescence.

Ciliates are important bacterial predators, mainly in nutrient-rich freshwater environments (e.g. Šimek e*t al.,* 1998, 2000; Thouvenot *et al.,* 1999), and are generally considered one of the key agents ensuring nutrient recycling and transfer to other trophic levels (e.g. Laybourn-Parry, 1992). We have confirmed and quantified bacterivory in *T. utriculariae*, whose grazing rates were comparable to literature reports (Neurer and Cowles, 1995). Due to their abundance, the ciliates were able to turn over the entire bacterial standing stock in the trap fluid extremely fast: 4 – 5 times in 24 hours in the younger traps (Table 3). Turnover time increased markedly in the old traps, due to the large increase in bacterial numbers. Obviously, the turnover of microbial mass in the *U. reflexa* traps and the release of soluble mineral nutrients from microbial cells with their subsequent uptake by the plant from the trap fluid are, to a large extent, facilitated by protist predation on bacteria. *Utricularia* plants apparently depend on microbial activity for the supply of nutrients and thus the amount of bacterial mass produced in the prey-free traps likely is as important to the plant host as the amount of organic matter digested.

### Predatory bacteria

Protists are, however, not the only predators affecting bacterial populations in the traps of *Utricularia.* We have also detected the family Bacteriovoracaceae in the lumen, whose members are known for being obligatory predators of other, especially Gram-negative, bacteria. There have been few studies of the ecological roles of predatory bacteria. They are, however, present in diverse habitats, which indicates that, like viruses, they are important determinants of microbial community dynamics (Velicer and Mendes-Soares, 2009). In the *Utricularia* traps, they are relatively abundant (Supplementary Table 2a), metabolically active (Supplementary Table 4) and, therefore, likely to selectively influence the trap microbial community dynamics through enhanced mortality rates of particular bacterial species in this closed environment.

### Microbial activity

When considering the overall activity of trap-associated microbiota, our results confirm highly dynamic, rapidly growing communities of both prokaryotes and eukaryotes (Tables 4a, 4b). Expression profiles (Supplementary Table 4) indicate rapid growth, intensive protein metabolism, respiration, DNA synthesis, and motility. Notable is the high expression of bacterial UDP-glucose-6-dehydrogenase, which has been linked to the environmentally regulated biosynthesis of exopolysaccharides (Petit *et al*., 1995), or that of transaldolase, which is also one of the highly expressed proteins in the trap fluid and has been shown to be an important colonization factor favoring the establishment of bacteria in the gut (González-Rodríguez *et al*., 2012). This activity is likely linked to bacterial aggregation and attachment to organic particles, which is typical for the rumen environment and has been observed in *Utricularia* traps previously (Sirová *et al*., 2009).

The transcriptomic analysis also offers several clues indicating the ability of trap-associated microbes to degrade complex organic material of algal origin. Algal cell walls and other cellular structures are composed of various monosacharides, derived from glucose, linked into polymers (cellulose and hemicellulose). These monosaccharides also include *D -* galactose (Buchanan *et al.*, 2015). Three enzymes involved in galactose metabolism were among the prokaryotic genes most expressed in *Utricularia* traps (Table 1a, Supplementary Table 4): the α-and β-galactosidases, which catalyze the cleavage of the terminal *D -* galactosyl residues of plant and algal hemicelluloses and their activity is often associated with herbivore digestive systems (Yapi *et al.*, 2007; Lu *et al*., 2012), and UDP-glucose 4-epimerase, which performs the final step in the Leloir pathway catalyzing the reversible conversion of UDP-galactose to UDP-glucose. The high expression levels of these enzymes underscore the importance of microbial galactose metabolism in the traps and are a further indication that the trap microbes, especially bacteria, actively degrade complex algal material.

### Conclusions

We conclude that, in analogy to herbivores, the aquatic *Utricularia* plants support the development of a diverse and sophisticated microbial ecosystem inhabiting both their traps and their large external surfaces, which allows them to digest and utilize complex organic material of algal origin and thrive even at highly oligotrophic sites. Here, although metazoan prey is usually scarce and prey capture rates correspondingly low (Richards, 2001), the supply of algae growing in close proximity to the traps as part of the *Utricularia* periphytic “gardens” is continuous and abundant (Płachno *et al*., 2012). In analogy to the digestive tracts of herbivorous animals, feedstuffs enter the traps and are degraded to various extents by the microbial populations colonizing them. Like the gut/ruminal ecosystems, traps harbor a diverse, symbiotic population of bacteria, fungi, and ciliates that have adapted for survival under low oxygen supply, high cell densities, and ciliate predation. Moreover, they have evolved the capacity for efficient utilization of complex and often recalcitrant plant polymers, such as cellulose and hemicellulose. As such, the *Utricularia* – microbe systems represent unique biodiversity and activity hotspots within the nutrient-poor, dystrophic environments, in which they grow (Figure 5), and should be considered as synchronized, mutually dependent biological and ecological units in future research.

**Figure 5.**
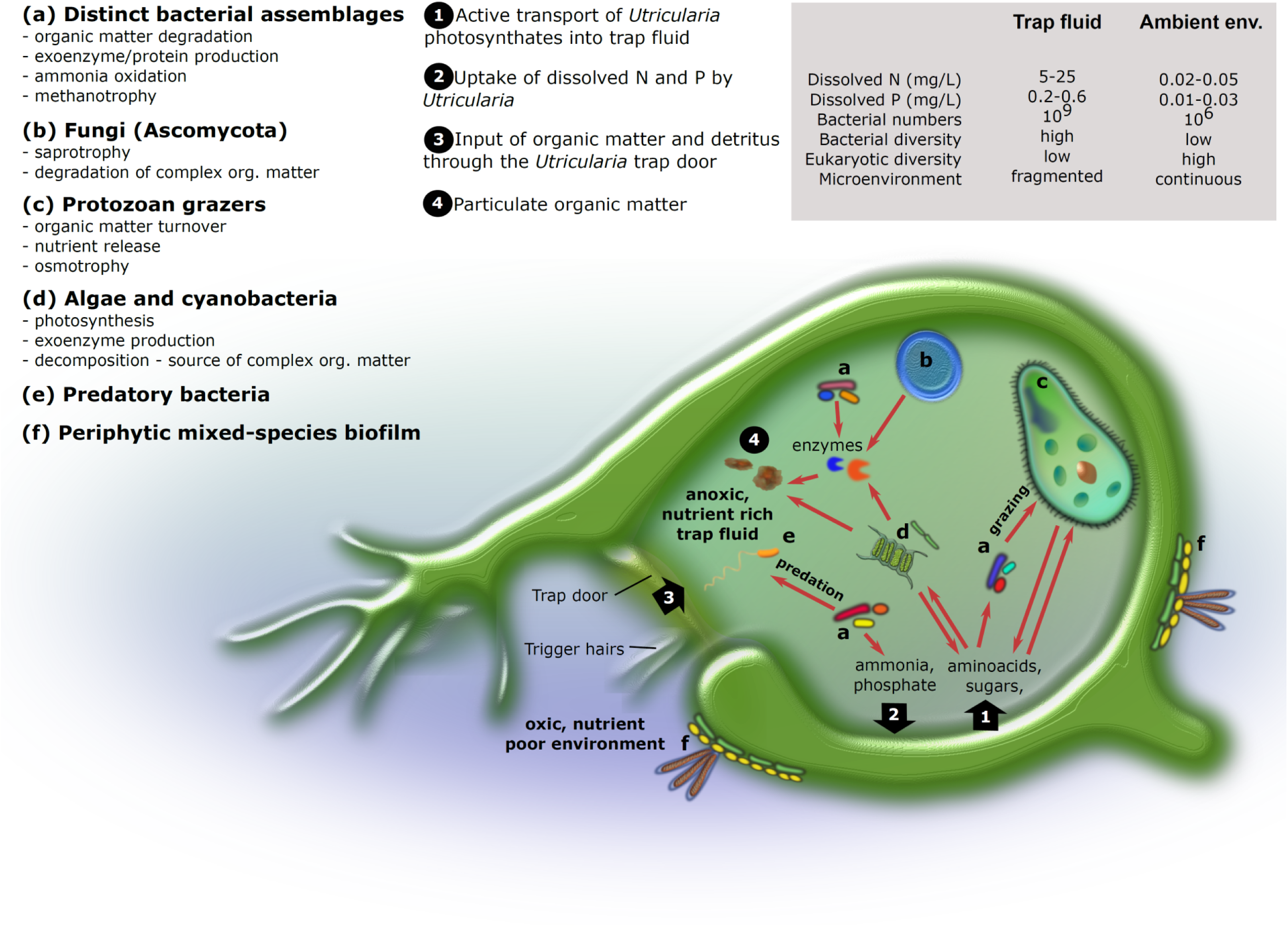
Conceptual representation of the *Utricularia* trap ecophysiology: main microbemicrobe and plant-microbe interactions are shown.

## Acknowledgements

We would like to acknowledge Dr. Helena Štorchová (Institute of Experimental Botany CAS) for her help and enthusiastic support of the project. This study was funded by the Czech Research Projects CSF P504/11/0783 (to LA) and CSF P504 – 17 – 10493S (to DS), and the Long-term research development project No. RVO 67985939 (to LA).

## References

Abraham JV, Butler RD, Sigee DC. (1997). Ciliate populations and metals in an activated-sludge plant. Water Res 31: 1103–1111.

Adamec L. (2007). Oxygen concentrations inside the traps of the carnivorous plants *Utricularia* and *Genlisea* (Lentibulariaceae). Ann Bot 100: 849–856.

Adamec L. (2011). Functional characteristics of traps of aquatic carnivorous *Utricularia* species. Aquat. Bot. 95: 226–233.

Adamec L, Sirová D, Vrba J. (2010a). Contrasting growth effects of prey capture in two aquatic carnivorous plant species. Fundam Appl Limnol / Arch fϋr Hydrobiol 176: 153–160.

Adamec L, Sirová D, Vrba J, Rejmánková E. (2010b). Enzyme production in the traps of aquatic *Utricularia* species. Biologia (Bratisl.) 65: 273–278.

Adlassnig W, Peroutka M, Lendl T. (2011). Traps of carnivorous pitcher plants as a habitat: Composition of the fluid, biodiversity and mutualistic activities. Ann Bot 107: 181–194.

Alcaraz LD, Martínez-Sánchez S, Torres I, Ibarra Laclette E, and Herrera-Estrella L. (2016). The metagenome of *Utricularia gibba*’s traps: Into the microbial input to a carnivorous plant. Plos One 11: e0148979.

Alkhalaf IA, Hϋbener T, Porembski S. (2009). Prey spectra of aquatic *Utricularia* species (Lentibulariaceae) in northeastern Germany: The role of planktonic algae. Flora Morphol Distrib Funct Ecol Plants 204: 700–708.

Attwood GT, Klieve A V., Ouwerkerk D, Patel BKC. (1998). Ammonia - hyperproducing bacteria from New Zealand ruminants. Appl Environ Microbiol 64: 1796–1804.

Barboza PS, Bennett a, Lignot J - H, Mackie RI, McWhorter TJ, Secor SM, et al. (2010). Digestive challenges for vertebrate animals: microbial diversity, cardiorespiratory coupling, and dietary specialization. Physiol Biochem Zool 83: 764–74.

Bárta J, Stone JD, Pech J, Sirová D, Adamec L, Campbell MA, et al. (2015). The transcriptome of *Utricularia vulgaris*, a rootless plant with minimalist genome, reveals extreme alternative splicing and only moderate sequence similarity with *Utricularia gibba*. BMC Plant Biol 15: 78.

Bates ST, Berg-Lyons D, Caporaso JG, Walters WA, Knight R, Fierer N. (2011). Examining the global distribution of dominant archaeal populations in soil. ISME J 5: 908–917.

Berg G, Grube M, Schloter M, Smalla K. (2014). Unraveling the plant microbiome: looking back and future perspectives. Front Microbiol 5: 148.

Bergmann GT, Bates ST, Eilers KG, Lauber CL, Caporaso JG, Walters WA, et al. (2011). The under - recognized dominance of Verrucomicrobia in soil bacterial communities. Soil Biol Biochem 43: 1450–1455.

Berry D, Widder S. (2014). Deciphering microbial interactions and detecting keystone species with co-occurrence networks. Front Microbiol 5: 219.

Borneman J, Hartin RJ. (2000). PCR primers that amplify fungal rRNA genes from environmental samples. Appl Environ Microbiol 66: 4356–60.

Borovec J, Sirová D, Adamec L. (2012). Light as a factor affecting the concentration of simple organics in the traps of aquatic carnivorous *Utricularia* species. Fundam Appl Limnol / Arch fϋr Hydrobiol 181: 159–166.

Buchanan BB, Gruissem W, Jones RL (Eds.). (2015). Biochemistry and molecular biology of plants, 2nd Edition. John Wiley and Sons, Ltd, Chichester, UK; pp. 45–90.

Caporaso JG, Kuczynski J, Stombaugh J, Bittinger K, Bushman FD, Costello EK, et al. (2010). QIIME allows analysis of high - throughput community sequencing data. Nat Methods 7: 335–336.

Caravieri FA, Ferreira AJ, Ferreira A, Clivati D, de Miranda VFO, Araújo WL. (2014). Bacterial community associated with traps of the carnivorous plants *Utricularia hydrocarpa* and *Genlisea filiformis*. Aquat Bot 116: 8–12.

Carretero-Paulet L, Librado P, Tien-Hao C, Ibarra-Laclette E, Herrera-Estrella L, Rozas J, Albert VA. (2015). High gene family turnover rates and gene space adaptation in the compact genome of the carnivorous plant Utricularia gibba. Mol Biol Evol 32: 1284 – 1295.

Cochran-Stafira DL, Von-Ende CN. (1998). Integrating bacteria into food webs: Studies with *Sarracenia purpurea* inquilines. Ecology 79: 880–898.

Darwin C. (1875). Insectivorous plants. New York: D Appleton and Company.

Edgar RC. (2010). Search and clustering orders of magnitude faster than BLAST. Bioinformatics 26:2460 – 2461.

Díaz-Olarte J, Valoyes-Valois V, Guisande C, Torres NN, González-Bermúdez A, SanabriaAranda L, et al.. (2007). Periphyton and phytoplankton associated with the tropical carnivorous plant *Utricularia foliosa*. Aquat Bot 87: 285–91.

Faust K, Sathirapongsasuti JF, Izard J, Segata N, Gevers D, Raes J, Huttenhower C. (2012). Microbial co-occurrence relationships in the human microbiome. PLoS Comput Biol 8: e1002606.

Friday LE. (1989). Rapid turnover of traps in *Utricularia vulgaris* L. Oecologia 80: 272–277.

González-Rodríguez I, Sánchez B, Ruiz L, Turroni F, Ventura M, Ruas-Madiedo P, Gueimonde M, Margolles A. (2012). Role of extracellular transaldolase from *Bifidobacterium bifidum* in mucin adhesion and aggregation. Appl Environ Microbiol 78: 3992–3998.

Gray SM, Akob DM, Green SJ, Kostka JE. (2012). The bacterial composition within the *Sarracenia purpurea* model system: local scale differences and the relationship with the other members of the food web. PLoS One 7: e50969.

Haas BJ, Papanicolaou A, Yassour M, Grabherr M, Blood PD, Bowden J, et al.. (2013). De novo transcript sequence reconstruction from RNA - seq using the Trinity platform for reference generation and analysis. Nat Protoc 8:1494–512.

Hackstein JHP, Van-Alen TA (1996). Fecal methanogens and vertebrate evolution. Evolution 50: 559–572.

Hales BA, Edwards C, Ritchie DA, Hall G, Pickup RW, Saunders JR. (1996). Isolation and identification of methanogen - specific DNA from blanket bog peat by PCR amplification and sequence analysis. Appl Environ Microb 62: 668–75.

Ibarra-Laclette E, Albert V, Pérez-Torres C, Zamudio-Hernández F, Ortega - Estrada MDJ, Herrera - Estrella A, et al. (2011). Transcriptomics and molecular evolutionary rate analysis of the bladderwort (Utricularia), a carnivorous plant with a minimal genome. BMC Plant Biol 11: 101.

James BR, and Wilson DB. (1996). Why are ruminal cellulolytic bacteria unable to digest cellulose at low pH? Journal of Dairy Science 79: 1503–1509.

Kolb S, Knief C, Stubner S, Conrad R. (2003). Quantitative detection of methanotrophs in soil by novel pmoA - targeted Real - Time PCR assays. Appl Environ Microbiol 69: 2423–2429.

Koopman MM, Fuselier DM, Hird S, Carstens BC. (2010). The carnivorous pale pitcher plant harbors diverse, distinct, and time - dependent bacterial communities. Appl Environ Microbiol 76: 1851–60.

Laybourn-Parry J. (1992). Protozoan plankton ecology. Chapman and Hall, London.

Leininger S, Urich T, Schloter M, Schwark L, Qi J, Nicol GW, et al. (2006). Archaea predominate among ammonia - oxidizing prokaryotes in soils. Nature 442: 806–809.

Liu Z, Lozupone C, Hamady M, Bushman FD, Knight R. (2007). Short pyrosequencing reads suffice for accurate microbial community analysis. Nucleic Acids Res 35. e - pub ahead of print, doi:10.1093 / nar / gkm541.

Lu HP, Wang YB, Huang SW, Lin CY, Wu M, Hsieh CH, Yu HT. (2012). Metagenomic analysis reveals a functional signature for biomass degradation by cecal microbiota in the leaf - eating flying squirrel (*Petaurista alborufus lena*). BMC Genomics 13: 466.

McAllister TA, Bae HD, Jones GA, Cheng KJ. (1994). Microbial attachment and feed digestion in the rumen. J Anim Sci 72: 3004–3018.

McFall-Ngai M, Hadfield MG, Bosch TCG, Carey H V, Domazet - LoŠo T, Douglas AE, et al. (2013). Animals in a bacterial world, a new imperative for the life sciences. Proc Natl Acad Sci U S A 110: 3229–36.

Muyzer G, de-Waal EC, Uitterlinden AG. (1993). Profiling of complex microbial populations by denaturing gradient gel electrophoresis analysis of polymerase chain reaction - amplified genes coding for 16S rRNA. Appl Environ Microbiol 59:695–700.

Nalepa C A., Bignell DE, Bandi C. (2001). Detritivory, coprophagy, and the evolution of digestive mutualisms in Dictyoptera. Insectes Soc 48: 194–201.

Neuer S, Cowles TJ. (1995). Comparative size - specific grazing rates in field populations of ciliates and dinoflagellates. Mar Ecol Prog Ser 125: 259–267.

Peroutka M, Adlassnig W, Volgger M, Lendl T, Url WG, Lichtscheidl IK. (2008). *Utricularia*: A vegetarian carnivorous plant? Agae as prey of bladderwort in oligotrophic bogs. Plant Ecol 199: 153–162.

Petit C, Rigg GP, Pazzani C, Smith A, Sieberth V, Stevens M, Boulnois G, Jann K, Roberts IS. (1995). Region 2 of the *Escherichia coli* K5 capsule gene cluster encoding proteins for the biosynthesis of the K5 polysaccharide. Mol Microbiol 17: 611–620.

Pitsch G, Adamec L, Dirren S, Nitsche F, Šimek K, Sirová D, Posch T. (2016). The green *Tetrahymena utriculariae* n. sp. (Ciliophora, Oligohymenophorea) with its endosymbiotic algae (*Micractinium* sp.), living in the feeding traps of a carnivorous aquatic plant. J Eukaryot Microbiol: doi:10.1111 / jeu.12369.

Płachno BJ, Łukaszek M, Wołowski K, Adamec L, Stolarczyk P. (2012). Aging of *Utricularia* traps and variability of microorganisms associated with that microhabitat. Aquat Bot 97: 44–8.

Płachno BJ, Wołowski K, Fleischmann A, Lowrie A, Łukaszek M. (2014). Algae and prey associated with traps of the Australian carnivorous plant *Utricularia volubilis* (Lentibulariaceae: *Utricularia* subgenus *Polypompholyx*) in natural habitat and in cultivation. Austral J Bot 62: 528–36.

Repeta DJ, Ferrón S, Sosa OA, Johnson CG, Repeta LD, Acker M, DeLong EF, Karl DM: (2016). Marine methane paradox explained by bacterial degradation of dissolved organic matter. Nature Geoscience 9: 884–887.

Richards JH. (2001). Bladder function in *Utricularia purpurea* (Lentibulariaceae): Is carnivory important? Am J Bot 88: 170–176.

Rinke C, Schwientek P, Sczyrba A, Ivanova NN, Anderson IJ, Cheng J - F, et al. (2013). Insights into the phylogeny and coding potential of microbial dark matter. Nature 499: 431–437.

Russell JB, Wilson DB. (1996). Why are ruminal cellulolytic bacteria unable to digest cellulose at low pH? J Dairy Sci 79:1503–9.

Russell JB, Muck RE, Weimer PJ. (2009). Quantitative analysis of cellulose degradation and growth of cellulolytic bacteria in the rumen. FEMS Microbiol Ecol 67: 183–197.

Shannon P, Markiel A, Ozier O, Baliga NS, Wang JT, Ramage D, Amin N, Schwikowski B, Ideker T. (2003). Cytoscape: a software environment for integrated models of biomolecular interaction networks. Genome Res 13: 2498–2504.

Shimkets L, Dworkin M, Reichenbach H. (2006). The myxobacteria. In: Dworkin M, Falkow S, Rosenberg E, Schleifer KH, Stackebrandt E (eds): The prokaryotes, vol 7, 3rd edn. Springer, Berlin, pp. 31–115.

Silva SR, Diaz YCA, Penha HA, Pinheiro DG, Fernandes CC, Miranda VFO, Michael TP, Varani AM. (2016). The chloroplast genome of *Utricularia reniformis* sheds light on the evolution of the ndh gene complex of terrestrial carnivorous plants from the Lentibulariaceae family. Plos One 11: e0165176.

Šimek K, Bobková J, Macek M, Jiří N, Psenner R. (1995). Ciliate grazing on picoplankton in a eutrophic reservoir during the summer phytoplankton maximum: A study at the species and community level. Limnol Oceanogr 40: 1077–1090.

Šimek K, Armengol J, Comerma M, Garcia JC, Chrzanowski TH, Macek M, Nedoma J, StraŠkrabová V. (1998). Characteristics of protistan control of bacterial production in three reservoirs of different trophy. Internat Rev Hydrobiol 83: 485–494.

Šimek K, Jϋrgens K, Nedoma J, Comerma M, Armengol J. (2000). Ecological role and bacterial grazing of *Halteria* spp.: Small freshwater oligotrichs as dominant pelagic ciliate bacterivores. Aquat Microb Ecol 22: 43–56.

Šimek K, Pernthaler J, Weinbauer MG, Hornák K, Dolan JR, Nedoma J, et al. (2001). Changes in bacterial community cmposition and dynamics and viral mortality rates associated with enhanced flagellate grazing in a mesoeutrophic reservoir. Appl Environ Microbiol 67: 2723–2733.

Šimek K, Pitsch G, Salcher MM, Sirová D, Shabarova T, Adamec L. et al.. (2017). Ecological traits of the algae - bearing *Tetrahymena utriculariae* (Ciliophora) from traps of the aquatic carnivorous plant *Utricularia reflexa*. J Eukaryot Microbiol: doi:10.1111 / jeu.12368.

Sirová D, Adamec L, Vrba J. (2003). Enzymatic activities in traps of four aquatic species of the carnivorous genus Utricularia. New Phytol 159: 669–675.

Sirová D, Borovec J, černá B, Rejmánková E, Adamec L, Vrba J. (2009). Microbial community development in the traps of aquatic *Utricularia* species. Aquat Bot 90: 129–136.

Sirová D, Borovec J, Šantrůčková H, Šantrůček J, Vrba J, Adamec L. (2010). *Utricularia* carnivory revisited: Plants supply photosynthetic carbon to traps. J Exp Bot 61: 99–103.

Sirová D, Borovec J, Picek T, Adamec L, Nedbalová L, Vrba J. (2011). Ecological implications of organic carbon dynamics in the traps of aquatic carnivorous *Utricularia* plants. Funct Plant Biol 38: 583–593.

Sirová D, Šantrůček J, Adamec L, Bárta J, Borovec J, Pech J, et al. (2014). Dinitrogen fixation associated with shoots of aquatic carnivorous plants: Is it ecologically important? Ann Bot 114: 125–133.

Slobodkin A. 2014. The Family Peptostreptococcaceae. In: Rosenberg E (Ed.): The Prokaryotes: Firmicutes and Tenericutes, 4^th^ Edition. Springer - Verlag, Berlin, Heidelberg, Germany. pp. 291–302.

Towne G, Nagaraja TG, Brandt RT, Kemp KE. (1990). Ruminal ciliated protozoa in cattle fed finishing diets with or without supplemental fat. J Animal Sci 68:21509–2155.

Thouvenot A, Richardot M, Debroas D, Devaux J. (1999). Bacterivory of metazooplankton, ciliates and flagellates in a newly - flooded reservoir. J Plankton Res 21:1659–1679.

Urich T, Lanzen A, Qi J, Huson DH, Schleper C, Schuster SC. (2008). Simultaneous assessment of soil microbial community structure and function through analysis of the meta - transcriptome. Plos One 3: e2527.

Velicer GJ, Mendes-Soares H. (2009). Bacterial predators. Curr Biol 19:pR55–R56.

Yapi D, Yapi A, Gnakri D, Niamke SL, Kouame LP. (2009). Purification and biochemical characterization of a specific β - glucosidase from the digestive fluid of larvae of the palm weevil, Rhynchophorus palmarum. J Insect Sci 9: 4.

Zerbino DR, Birney E. (2008). Velvet: algorithms for de novo short read assembly using de Bruijn graphs. Genome Res 18: 821–829.

